# A cortical mechanism linking saliency detection and motor reactivity in rhesus monkeys

**DOI:** 10.1101/2023.01.31.526437

**Authors:** Giacomo Novembre, Irene Lacal, Diego Benusiglio, Eros Quarta, Andrea Schito, Stefano Grasso, Ludovica Caratelli, Roberto Caminiti, Alexandra Battaglia Mayer, Gian Domenico Iannetti

## Abstract

Sudden and surprising sensory events trigger neural processes that swiftly adjust behavior. To study the phylogenesis and the mechanism of this phenomenon, we trained two rhesus monkeys to keep a cursor inside a visual target by exerting force on an isometric joystick. We examined the effect of surprising auditory stimuli on exerted force, scalp electroencephalographic (EEG) activity, and local field potentials (LFP) recorded from the dorso-lateral prefrontal cortex. Auditory stimuli elicited (1) a biphasic modulation of isometric force: a transient decrease followed by a corrective tonic increase, and (2) EEG and LFP deflections dominated by two large negative-positive waves (N70 and P130). The EEG potential was maximal at the scalp vertex, in all respects similar to the human ‘vertex potential’. Electrocortical potentials and force were tightly coupled: the P130 amplitude predicted the magnitude of the corrective force increase, particularly in the EEG electrodes contralateral to the limb exerting force, and in the LFP recorded from deep rather than superficial cortical layers, suggesting a direct effect of the vertex potential on the motor output determining the behavior. These results disclose a phylogenetically-preserved cortico-motor mechanism supporting adaptive behavior in response to salient sensory events.

## Introduction

Survival in the natural world depends heavily on an animal’s capacity to identify threats or affordances, and to adapt ongoing behavior accordingly. Coping with the ever-changing nature of the surrounding environment, the human brain has developed the capacity to initiate rapid and adaptive behavioral responses to salient sensory stimuli, with none or scarce influence of volition. We recently referred to this as Reactive Adaptive Behavior (RAB): salient sensory stimuli elicit fast involuntary behavioral responses that are, however, flexible on the basis of the current environmental context (1).

There are multiple examples of RAB in the literature. One is cortico-muscular resonance (CMR) (2,3). This consists of a series of fast modulations of muscular activity evoked by sudden and unexpected sensory stimuli. The CMR has only been studied in humans so far: Participants are asked to exert a constant isometric force on a transducer held between the index finger and the thumb, while fast-rising and task-irrelevant unexpected stimuli are delivered (2,3). The stimuli evoke an initial force decrease (*d1*, peaking ~100 ms post-stimulus) followed by two consecutive force increases (*i1*, peaking at ~250 ms; *i2*, starting ~300-350 ms and lasting for ~2 s). These force modulations are tightly coupled to a large electro-cortical response elicited by the same stimuli evoking the CMR. This event-related EEG potential is dominated by two large negative-positive waves maximal at the scalp vertex and it is therefore named ‘vertex potential’ (4–6). Interestingly, the trial-by-trial amplitude of the vertex potential predicts the magnitude of the force modulations, particularly the two force increases (3).

The CMR falls within the definition of RAB. First, CMR is elicited in response to salient stimuli in an automatic and unconscious manner, i.e. participants are unaware of producing a response. Second, CMR is adaptive, e.g. the amplitude of the force modulations is enhanced if the eliciting stimulus has high behavioral relevance (1).

The CMR, and RABs in general, might be a fundamental backbone of animal survival. As such, one would guess that these behavioral responses are well conserved phylogenetically. Yet, whether CMR is also observable in other species besides humans is currently unknown. Nevertheless, other RABs besides CMR, such as stimulus-locked responses (7–10), online motor corrections (11–14) or action stopping (15,16), have been identified in both humans and non-human primates. This suggests that CMR might also be observable in non-human primates.

Therefore, the first aim of the current study was to investigate whether the CMR is also observable in non-human primates, specifically rhesus monkeys (*Macaca mulatta*). To do so, across two Experiments, we exploited a well-established behavioral task that requires monkeys to control the position and motion of a (visually-presented) cursor on a screen using a hand-held force-sensitive isometric joystick (Fig. 1a) (17,18). Animals were trained to hold the cursor inside a central target, an action that implied the production of a constant and fine-grained force, while isolated fast-rising and task-irrelevant auditory stimuli were presented on a minority of the trials (Beep Trials). This permitted us to examine the effect of such sudden and unexpected stimuli on the isometric force exerted by the monkeys, i.e. to test whether the CMR is also observable in non-human primates.

**Figure 1.**
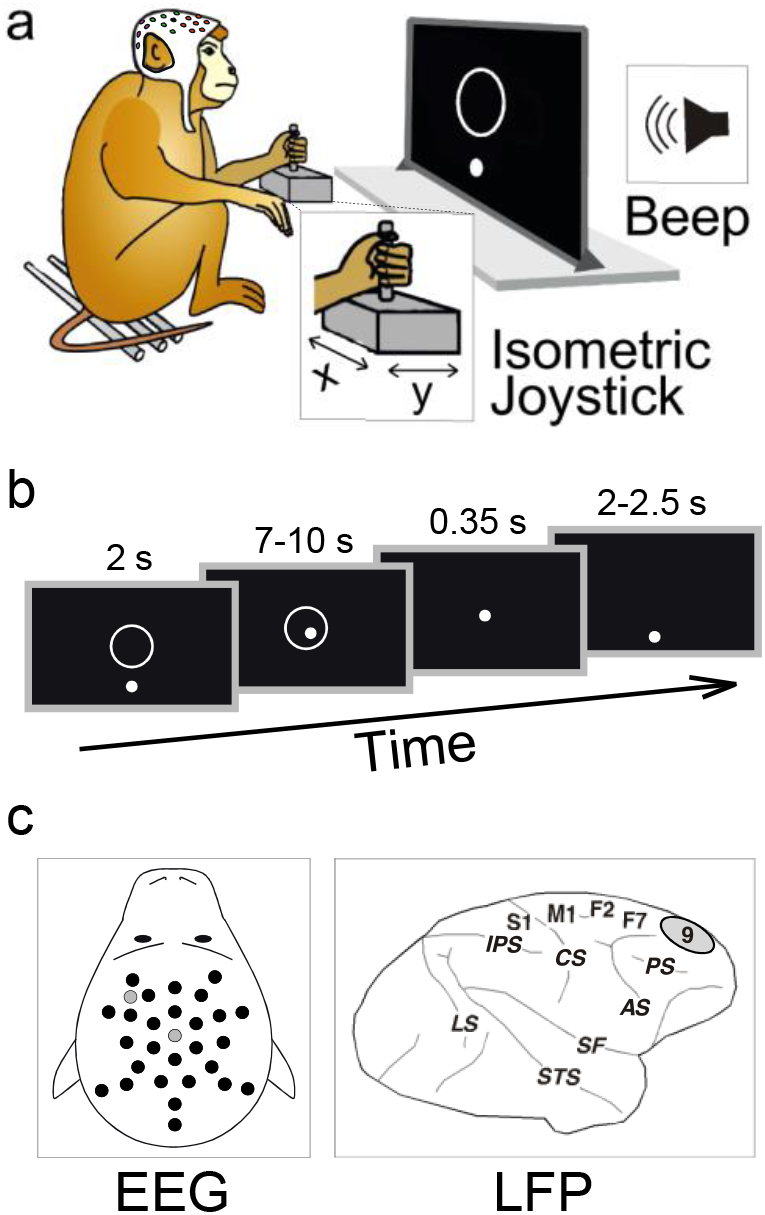
Experimental materials and methods. **(A) Experimental paradigm.** Two macaques were trained to exert a force on an isometric joystick using the left hand. The force applied on the x and y axes of the joystick was used to control the position of a cursor moving on a monitor. During a period of static force application, sudden task-irrelevant auditory stimuli were delivered through a beeper placed behind the monitor. **(B) Task timeline**. The task begun with presentation of the target on the center of the screen. Monkeys had 2 seconds to bring the cursor (white dot) inside the target (white circle) and were required to hold the cursor there for a variable time interval (ranging between 7 and 10 seconds). In 33% of the trials, auditory stimuli were unexpectedly delivered during this interval. If the cursor remained inside the target, the trial was considered successful, and a liquid reward was given. Trials were separated by a 2-2.5 second (jittered) interval (during which monkeys were not required to exert force and therefore the cursor was likely to be back to the start position). **(C) EEG and LFP recording.** In Experiment 1, we recorded EEG signals using 29 active electrodes (black dots) and 2 “zero-reference” electrodes (grey dots), mounted on custom-made EEG caps tailored to fit each animal’s head. In Experiment 2, local field potentials (LFP) were recorded from the right dorso-lateral prefrontal cortex (Brodmann area 9), through a 5-channel multiple-electrode array system for extracellular recording (see also Supplementary Figure 1).

The second aim of this study was to investigate the neurophysiological origins of CMR in monkeys. In Experiment 1, based on the previous demonstration of a tight coupling between saliency-related vertex potentials and CMR, we used 29 active electrodes to record electroencephalographic (EEG) activity in awake monkeys performing the task described above (Fig. 1a,c). We examined event-related EEG potentials elicited by the salient stimuli and their relationship with the CMR. In Experiment 2 we repeated the above procedure while replacing EEG with intracortical recordings of local field potentials (LFP) from the right dorso-lateral prefrontal cortex (dlPFC, putatively Brodmann Area 9, BA9) – a cortical area that has been shown to be involved in hand force control in both human (19,20) and non-human primates (21). Notably, one study demonstrated that BA9 lesioning leads to a bilateral impairment of fine hand force control, leaving general motor behavior intact (21).

Together, the EEG and LFP recordings allowed us to confirm the cortical origin of the CMR. Furthermore, the LFP recordings allowed us to compare the effect of responses measured at different cortical depths. This latter notion might shed light upon the specific circuits through which BA9 might contribute to the CMR.

## Results

### Stimulus-induced Force modulations (Experiment 1)

In both monkeys, auditory stimuli elicited a consistent biphasic modulation of the overall force magnitude (F; Fig. 2a third row): an initial force decrease was followed by a force increase. T-tests comparing the exerted force across Beep and No-Beep trials (i.e. trials during which there was not auditory stimulus, see Methods) confirmed the across-trial consistency of this observation, in each animal. To assist interpretability of these modulations with respect to their human equivalents, the initial force decrease and the following increase will be hereafter referred to as *d1* and *i2*, respectively.

**Figure 2.**
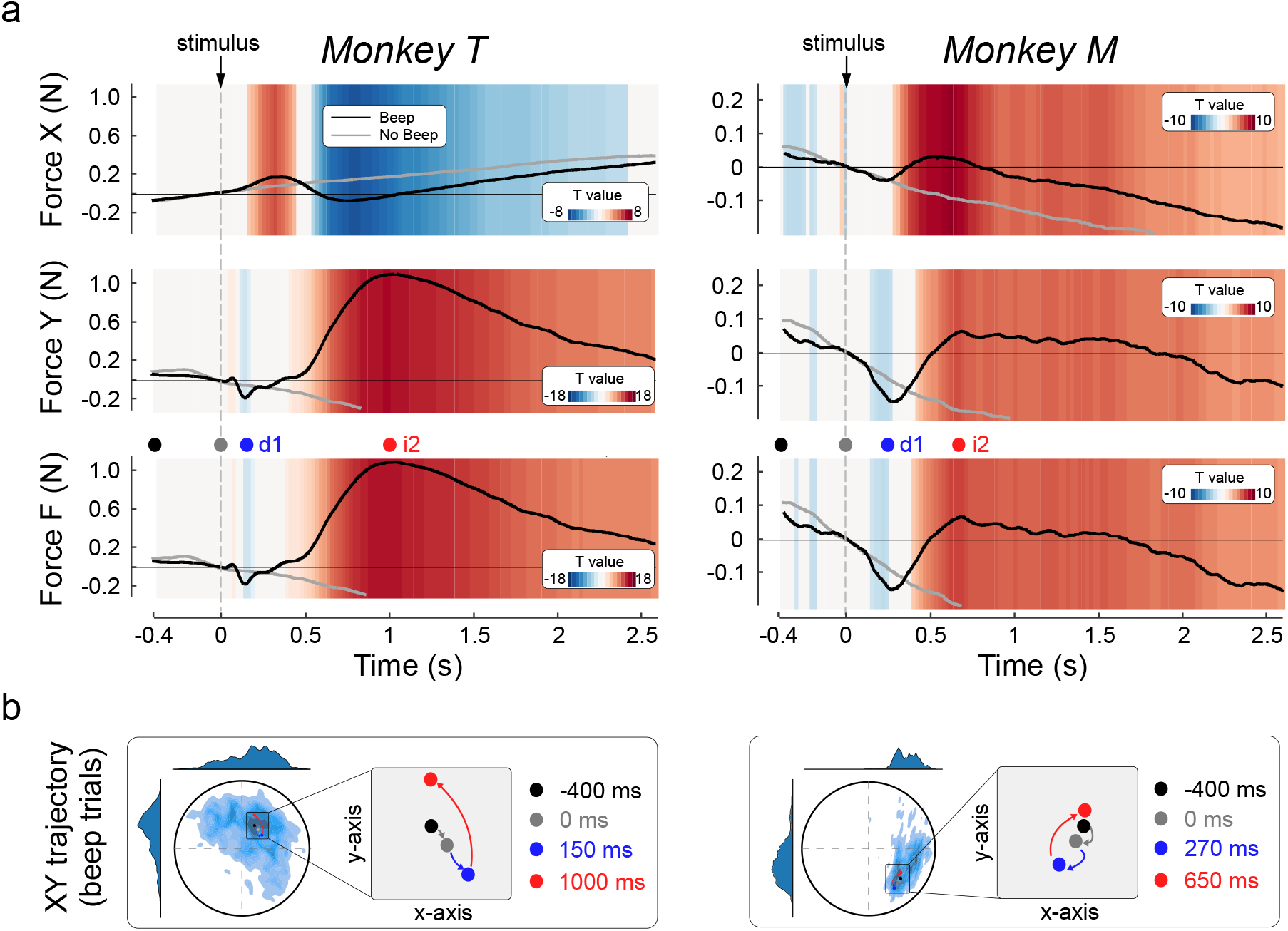
Stimulus-induced Force modulations. **A**: Stimulus-induced modulations of force magnitude over the x (first row) and y (second row) axes. A composite index of force (F), representing the overall force magnitude regardless of its x-y directionality, is displayed in the third row. The coloured background represents t values yielded after comparing Beep (black line) and No-Beep (grey line) trials. **B**: Illustrative representation of the position of the cursor (dot) with respect to the target (circle) over time, at four different time points: baseline onset (black), stimulus presentation (grey), peak of force decrease (blue) and peak of force increase (red). The density maps represent all positions held by the dot over the course of all trials.

The latency of the force modulation, particularly the initial *d1*, was slightly different across animals. In monkey T *d1* peaked at ~150 ms, while in monkey M it peaked at ~270 ms post-stimulus. In contrast, the subsequent *i2* was more similar across animals: it began ~400-450 ms post-stimulus and lasted nearly the whole trial duration. Notably, and paralleling human observations (3,22) in both animals *d1* had a more transient character, while *i2* was more tonic.

Examining the simultaneous modulations of force separately on the x and y axes (Fig. 2a, first and second row), we reconstructed the average cursor trajectory before and after the presentation of the auditory stimulus (Fig. 2b). In both monkeys, prior to stimulus presentation the cursor slowly drifted towards the bottom of the screen (black arrow, Fig. 2b). Bearing in mind that a force resulting in an upward movement on the y axis had to be exerted to keep the cursor inside the target, this observation is consistent with the well-known fatigue effect in isometric force tasks [which we and others also observed in humans; (3,23)]. Immediately after stimulus onset, the first force decrease (*d1*) resulted in a transient enhancement of the above-described pre-stimulus drift (blue arrow, Fig. 2b). The subsequent force rebound (*i2*) moved the cursor in the opposite direction, bringing it above the pre-stimulus position (red arrow, Fig. 2b).

Comparing the direction of these motion trajectories across monkeys, we noticed that they were consistent along the vertical y axis, but somehow different along the horizontal x axis: in monkey T the cursor drifted towards the right side of the monitor, while in monkey M it drifted to the left. This difference is possibly explained by the different position of each monkey relative to the monitor (slightly on the right-side of monkey M and on the left-side of monkey T; see *experimental setup*). Thus, the different drifting along the x axis might be trivially explained by the different hand and arm posture of the two animals.

### Stimulus-induced EEG modulations (Experiment 1)

The EEG modulation elicited by the auditory stimuli is displayed on Figure 3. The modulation of EEG voltage consisted of a triphasic pattern including an early positivity (P30), a negativity (N70) and a second longer lasting positivity (P130). The negative-positive N70 and P130 complex constitutes the well-known vertex potential that can be measured in human and non-human primates (4,5,24–27)

**Figure 3.**
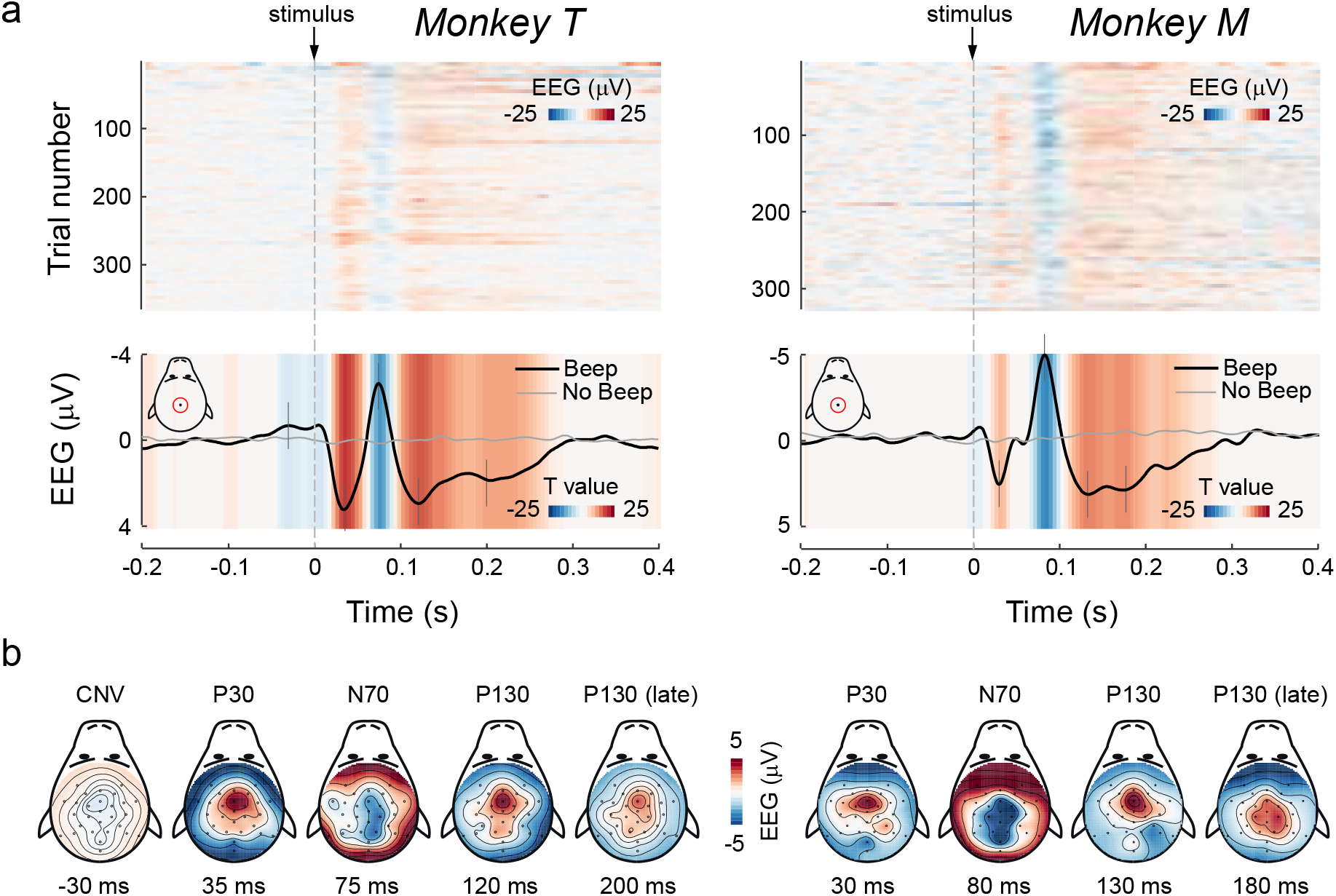
Stimulus-induced EEG modulations. **A (top)**: Single-trial modulations (at electrode Cz). Trials are sorted by their order of occurrence. The coloured background represents amplitude. **(bottom)**: across-trial averages of EEG modulations (at electrode Cz). The coloured background represents t values yielded after comparing Beep (black line) and No-Beep (grey line) trials. **B**: EEG topographies of the main modulations. Time points of each topography are marked with vertical grey lines crossing the EEG average waveform (shown in panel A, bottom).

Both latencies and topographies of these EEG modulations were remarkably consistent across animals. Specifically, the P30, which had central and frontal distribution, peaked at 35 and 30 ms post-stimulus in T and M, respectively. The N70 had broader and more posterior distribution over the scalp, and it peaked at 75 and 80 ms in T and M, respectively. Finally, the early part of the P130 exhibited a central-frontal topography, peaking at 120 and 130 ms in T and M, respectively. Notably, the P130 lasted longer than the previous P30 and N70, and its initial frontal topography changed slightly throughout time, to become more widespread and centrally distributed ~180-200 ms post stimulus, particularly in monkey M (Fig. 3b). The t-tests comparing the EEG voltages associated to Beep and No-Beep trials confirmed the high across-trial consistency of all of the described components (Fig. 3a, bottom).

Notably, monkey T exhibited a mild slow-rising negativity anticipating the stimulus. This component is most likely a contingent negative variation (CNV) (28,29).

### Trial-by-trial correlation between force and EEG modulations (Experiment 1)

The trial-by-trial correlation between force and EEG modulations revealed several interesting relationships, which are outlined in Figure 4.

**Figure 4.**
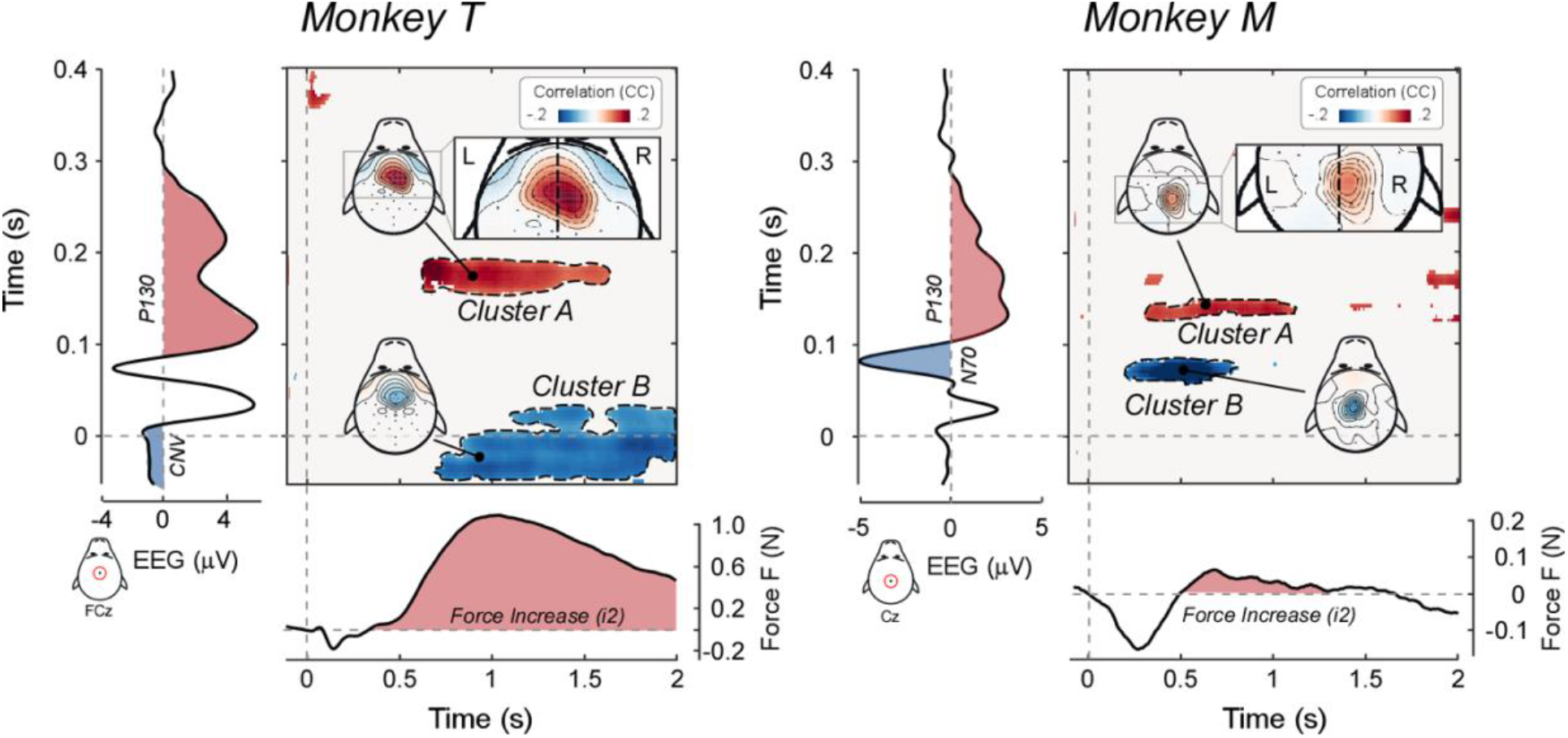
Trial-by-trial correlation between force and EEG modulations. Trial-by-trial correlations between stimulus-induced force and EEG modulations. Bidimensional plots represent the significant trial-by-trial correlation coefficients (cluster-corrected Spearman’s r) between EEG and force, for all possible pairs of time points, at electrode Cz. The topographies of the main correlation clusters are also plotted. The EEG timeseries (plotted vertically) and the force timeseries (plotted horizontally) are shown to assist interpretability of the correlations. Note that the correlation between the EEG positive wave (P130) and the force increase (i2) is slightly lateralized towards the right scalp regions, i. e. contralateral to the (left) arm that exerted the force.

First, both monkeys exhibited a robust correlation between the P130 EEG wave and the force increase *i2* (cluster A, Fig. 4). This implies that trials in which the P130 had large amplitude were also associated with a large force increase. It is also important to examine *where* across the scalp this correlation occurred [i.e. where trial-by-trial fluctuations of EEG amplitude were more strongly coupled with fluctuations of *i2* magnitude, see (3)]. In both animals, this correlation was stronger over the right hemisphere, i.e. contralaterally to the (left) hand exerting the force (Fig. 4, inset). This pattern is consistent with an effect of the stimulus-induced EEG modulation (i.e. the vertex potential) on cortical motor regions contralateral to the hand performing the task. Remarkably, both the correlation between the positive vertex potential and the *i2*, and the topography of such correlation were strikingly similar to what we previously observed in humans (2,3).

We also observed two additional relationships between EEG and force modulations that, however, were not consistent across the two animals (clusters B, Fig. 4). First, in monkey M, the amplitude of the N70 correlated negatively with the magnitude of the force increase following *d1* (i.e. with the ascending branch of *d1* and the initial part of *i2*) – another result that parallels what we observed in humans (3). Second, in monkey T, the amplitude of the CNV correlated negatively with the magnitude of *i2*.

### Trial-by-trial correlation between force and LFP modulations

Experiment 2 revealed a pattern of force modulation broadly similar to the one observed in Experiment 1 (compare Figs. 2 and 5). In both monkeys, auditory stimuli elicited modulations of the overall force magnitude (F) in a biphasic pattern composed of an initial force decrease (*d1*) followed by a force increase (*i2*). In monkey M, *d1* peaked at 148 ms post-stimulus, while *i2* peaked at 359 ms post-stimulus. In monkey T, *d1* showed a double peak (at 163 and 409 ms post-stimulus), due to an additional force increase peaking at 281 ms post-stimulus. The late force increase *i2* started ~400 ms post-stimulus and peaked >1 s post-stimulus. The morphology of these force responses, specifically that of the *i2*, was similar to that described above (Figs. 2 and 5).

**Figure 5.**
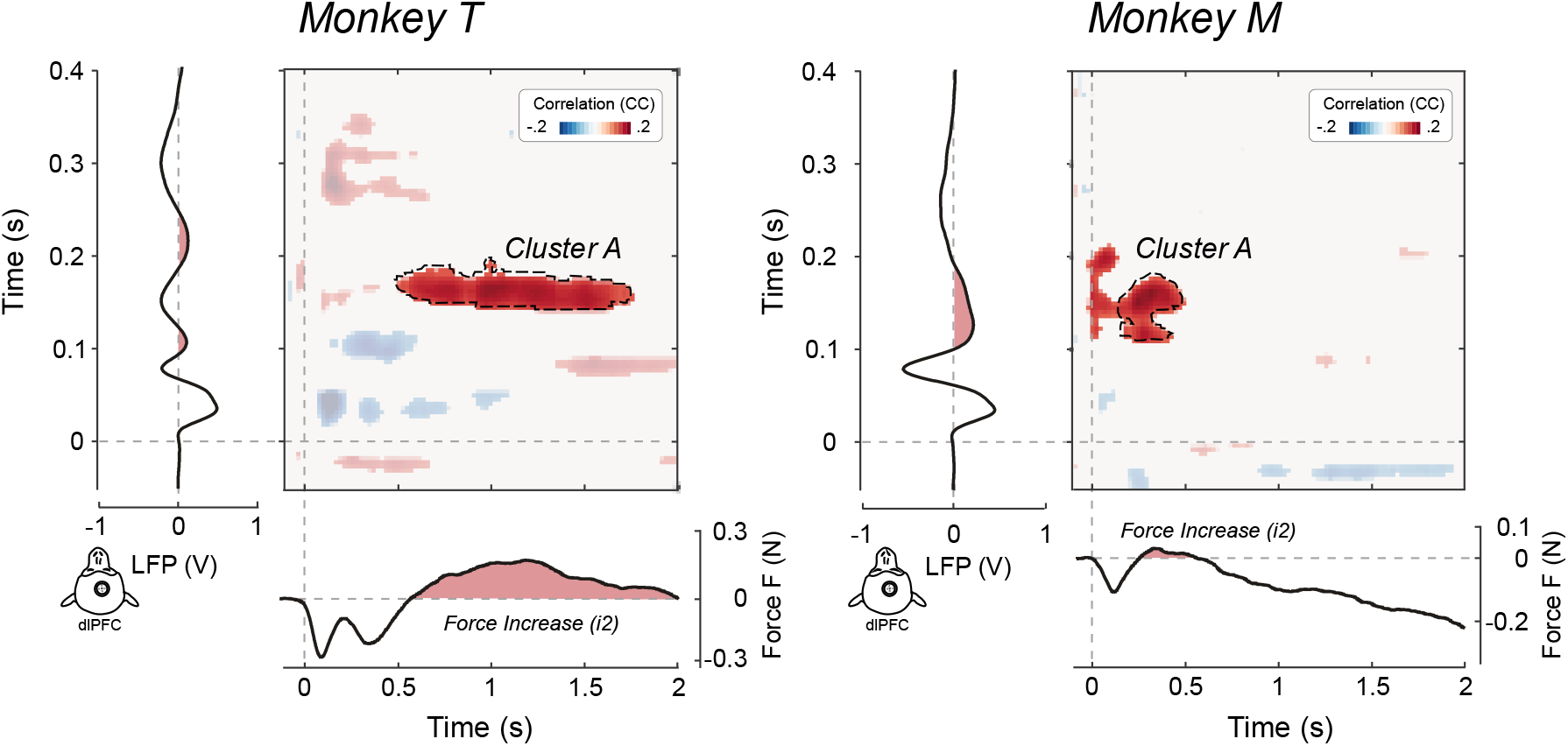
Trial-by-trial correlation between force and LFP modulations. Trial-by-trial correlations between stimulus-induced force and LFP modulations (recorded from the dorso-lateral Prefrontal Cortex, dlPFC). The bidimensional plots represent the significant trial-by-trial correlation coefficients (cluster-corrected Spearman’s r) between LFP and force, for all possible pairs of time points (pooling all ‘active’ electrodes). The LFP timeseries (plotted vertically) and the force timeseries (plotted horizontally) are shown to assist interpretability of the correlations. The correlation between the LFP positive wave (equivalent to the EEG P130) and the force increase (i2) is highlighted.

The auditory stimuli also elicited LFP modulations markedly similar to the EEG modulations described above (compare Experiment 1 and 2, Fig. 3 and 5). Specifically, these modulations entailed a triphasic pattern consisting of an early positivity (36 ms post-stimulus in both M and T), a negativity (78-79 ms in both M and T) and a second longer lasting positivity. In monkey M, this last positivity was very similar to what observed in Experiment 1 and peaked at 127 ms post-stimulus. In monkey T, this positive component appeared to be split into two halves (peaking at 106 and 215 ms post-stimulus, respectively), due to an additional negative deflection peaking at 151 ms post-stimulus. Looking more closely to the results from Experiment 1, this negativity embedded within the last P wave was also present in the EEG data (Figs. 3 and 4, left), although less clearly than in the LFP data (Fig. 5).

Most compellingly, the correlation between LFP and Force data was extremely similar to that observed between EEG and Force (compare Figs. 4 and 5). Specifically, the late positivity evoked by the auditory stimulus correlated, on a trial-by-trial level, with the late force increase *i2*, in both animals (Cluster A, Fig. 5). When we looked at this correlation as a function of cortical depth, i.e. considering selectively deep and superficial recording sites, we found that the correlation between LFP and Force was clearer for deep electrodes (Supplementary Fig. 2).

## Discussion

The current study was designed to address two questions. First, whether the CMR – a multiphasic modulation of isometric force elicited by salient sensory stimuli – is observable in non-human primates, particularly in rhesus monkeys. Second, combining force recordings with simultaneous EEG (experiment 1) and LFP (experiment 2) recordings, we investigated the neurophysiological origins of such (monkey) CMR.

In the next sections, we first describe the CMR in the macaque and compare it with the CMR observed in humans. Next, we describe the EEG/LFP responses elicited by the same stimuli causing the CMR, with particular emphasis on the vertex potential. We conclude by discussing the cortical origin of the CMR, focusing on the coupling between the vertex potential and the CMR-like response observed in monkeys.

### Force modulations: CMR in rhesus monkeys?

In our previous work in humans, we showed that salient auditory stimuli evoke a complex modulation of constantly-applied isometric force, named CMR (2,3,30). The human CMR consists in an initial force decrease peaking at 100 ms post-stimulus (*d1*), followed by two force increases: one peaking at 250 ms post-stimulus (*i1*) and the other starting at ~350 ms and lasting for nearly 2 seconds (*i2*) (see Fig. 6). The results of the two current experiments – conduced with rhesus monkeys – are reminiscent of the previous human results but also exhibit some differences, which we now discuss in detail. We particularly focus on the force increase, firstly because of its reproducibility across animals and experiments in the current work, and secondly because of its correlation with electrocortical activity observed here and previously in humans.

**Figure 6.**
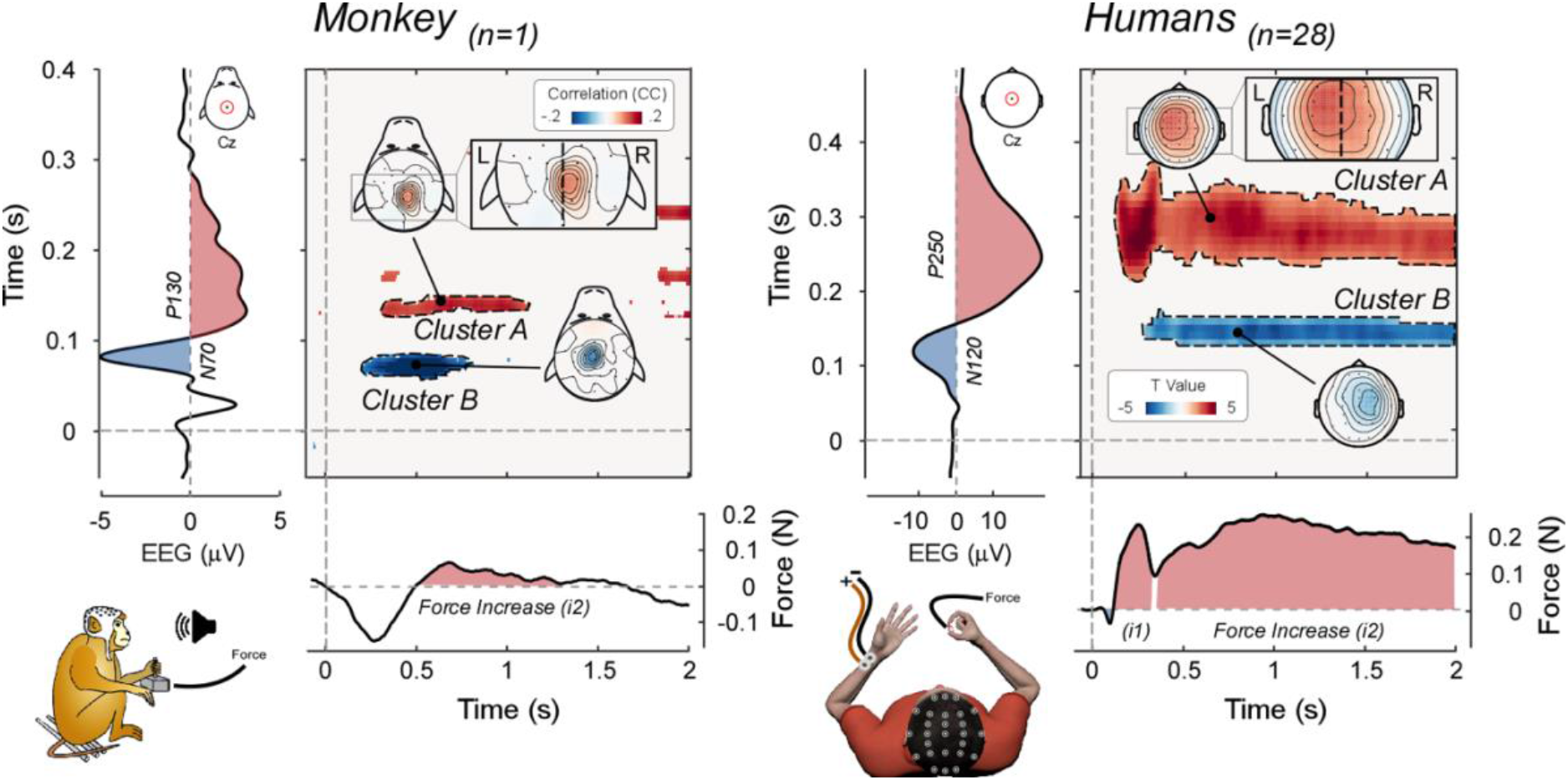
Comparison of stimulus-induced EEG-force correlations in monkeys and humans. Data are from the current study (Monkey M, left) and from (3) (28 human participants, right). The bidimensional plots represent the significant trial-by-trial correlation coefficients (cluster-corrected Spearman’s r [monkey, left], and t-values comparing participants’ Pearson’s r [human, right] between EEG and force for all possible pairs of time points, at electrode Cz (topographies of the highlighted clusters are plotted). The EEG timeseries (plotted vertically) and the force timeseries (plotted horizontally) are shown to assist interpretability of the correlations. Note that the correlation between the positive vertex wave (P130 in monkeys and P250 in humans) and the force increase (i2) is slightly lateralized towards the scalp regions contralateral to the hand that exerted the force (left hand in monkeys; right hand in human participants). Note that the two datasets were re-referenced differently, likely explaining the more focal (monkey) and global (human) topographies.

In both humans and monkeys, salient stimuli evoked an initial force decrease, followed by a force increase. However, while in humans we were able to distinguish two distinct force increases, this was mostly not the case in macaques (Fig. 6). This finding might be interpreted as evidence that only one increase is present in monkey (specifically *i2*, see below). Alternatively, bearing in mind that in the human experiments participants exerted an isometric force holding a transducer between the index and the thumb (i.e. a precision grip) while the monkeys held a joystick using their whole hand (i.e. power grip; see Figure 1 and 6), we could also hypothesize that the latter task does not lead to clearly distinguishable force increases.

The *i2* observed in monkeys started ~300-400 ms post-stimulus and was very tonic, lasting between 1s (monkey T) and 1.5 s (monkey M), remarkably similar to the human *i2*, which started ~350 ms post-stimulus and lasted for ~2 seconds (Fig. 6). The similarity in both latency and duration of these two force increases suggest that they might be homologous, hence we labelled the single force increase observed in monkeys as *i2*. Indeed, the human *i1* peaked at 250 ms post-stimulus and was much more transient (duration 200 ms, see Fig. 6), and, in the context of the current task and grip posture, it does not seem to have a consistent homologous in monkeys.

To appreciate the functional significance of such CMR response observed in the macaque, we reconstructed the cursor trajectory before and after the presentation of the auditory stimulus. We made several intriguing observations that might clarify the function of *d1* and *i2* (Fig. 2). The drift of the cursor towards the bottom of the monitor before stimulus presentation (see results for details) is likely to reflect the well-known fatigue effect often observed during isometric force tasks (3,23). Keeping this in mind, *d1* could be seen as a further transient reduction of the tonic corticospinal output that subserves task execution. Similarly, *i2* could be seen as a corrective rebound, bringing the cursor back to its original pre-stimulus position, possibly with some overshooting: note how the cursor position at the end of *i2* (red dots in the bottom panel of Figure 2) is higher than the cursor position 400 ms before stimulus onset (black dots in Figure 2). This is consistent with the idea that the CMR – and particularly its *i2* component – is both a reactive and an adaptive form of behavior (i.e. a RAB) (1).

### EEG/LFP modulations: the Vertex Potential in rhesus monkeys

The auditory stimuli evoked a number of transient modulations of the EEG (experiment 1) and LFP (experiment 2) recordings, which were very consistent both within and across animals (compare Figs. 3, 4 and 5). Specifically, we observed three post-stimulus modulations: an early positivity (P30), followed by a negativity (N70) and a last long-lasting positivity (P130). Such triphasic pattern has also been previously reported when auditory stimuli were presented to both rhesus (24,25) and squirrel monkeys (26,27).

In humans, auditory stimuli like those used here also evoke a triphasic pattern composed of an early positivity (P50), a negativity (N100) and a late positivity (P200). The latter two components, often labelled N1 and P2, constitute the widely-studied ‘vertex potential’, which is thought to index ‘surprise’ of the nervous system in response to salient (i.e. attention grabbing) stimuli, regardless of their sensory modality (4,5,30). The EEG/LFP modulations observed in monkeys are clearly reminiscent of the human vertex potential, although they exhibit slightly earlier latencies: the P30, the N70 and the P130 were 20, 30 and 70 ms shorter than their human homologues, respectively. These latency differences might be explained by the smaller length of the recruited fiber tracts of macaques vs humans, which implies shorter delays in information transfer (31–33).

Comparing EEG topographies across humans and macaques is usually difficult due to the different head shape and the presence of thick muscles surrounding the ears and neck of macaques (31). Despite this, the EEG topographies reported here, primarily centered over the scalp vertex, align extremely well with those commonly reported observed in humans (5,34) and with the few available reports of event-related potentials in monkeys (24,35). Possibly, these neat topographies benefitted from the fact that we capitalized on a combination of well-established and recently-developed EEG denoising algorithms, namely independent component analysis (ICA, REF) and artifact subspace reconstruction (ASR, (36,37)), which have rarely or never been applied to non-human primate EEG data. To the best of our knowledge, the presented results are one of the (if not the) most comprehensive characterizations of event-related potentials elicited by unexpected sensory events in non-human primates. This analytical development might fertilize other communities of scientists recording EEG from monkeys (or other non-human species), often not providing clear EEG topographical information (25,38–42).

### Cortical origin of the CMR

The second objective of the present study was to investigate the neural origin of the CMR. In humans, we demonstrated that the modulations of isometric force are tightly coupled to modulations of electro-cortical activity elicited by the same salient and unexpected stimuli evoking the CMR (2,3). Specifically, the trial-by-trial amplitude of both the negative (N100) and the positive (P200) waves composing the vertex potential elicited by the salient stimuli correlated with the magnitude of both force increases. The current experiment shows that a similar coupling is present in monkeys (Fig. 6).

In particular, we observed a robust trial-by-trial positive correlation between P130 (EEG/LFP; equivalent to the human P200) and *i2* (force), in both animals (clusters A, Fig. 4, 5). This correlation implies that trials associated with a large P130 amplitude were also associated with a large *i2*. Importantly, while the EEG topographical distribution of the P130 was symmetrical (Fig. 3b), the EEG scalp topography of this correlation was lateralized, and maximal on the hemisphere contralateral to the hand exerting the force (Fig. 4, insets). This discrepancy between voltage and correlation topographies, which corroborates what we previously observed in humans (3), is important, as it suggests that corticospinal projections originating in the frontal cortex contralateral to the limb performing the task might be modulated by the vertex potentials. This point is further supported by the LFP results, especially when bearing in mind that LFPs were recorded from the right dorsolateral prefrontal cortex, i.e. contralaterally to the limb performing the task (Fig. 1 and 5). Thus, the EEG/force and the LFP/force correlations observed in monkeys replicate and extend our previous human observations and provide stronger evidence that the cortical and muscular responses elicited by sudden and unexpected environmental events are strongly coupled. Together, these results imply that the vertex potential, and specifically its P wave, reflects a phylogenetically conserved system that responds to salient environmental events with a large cortical response that directly affects the muscles through a modulation of corticofugal output, and thereby prompts rapid and appropriate behavioral responses.

We also observed other correlations that were inconsistent across the two tested animals, although sometimes consistent with human results (clusters B, Fig. 4, Fig. 6). These additional correlations should therefore be interpreted with caution. In monkey M, the trial-by-trial variability of the N70 amplitude (homologous of the human N100) correlated negatively with the magnitude of *i2*: thus, a larger N70 predicted the subsequent occurrence of a stronger *i2*. This is also consistent with human results (3). In particular, the fact that we observed the N70-*i2* correlation only in one animal, and the P130-*i2* correlation in both animals might be consistent with the observation that, in humans, the N100-*i2* correlation was less robust (*p*=0.019) than the P200-i2 correlation (*p*<0.001). Furthermore, in monkey T, the amplitude of the CNV correlated negatively with the magnitude of *i2*. Given that (1) this result was unexpected, (2) observed only in one animal (and not in humans), and (3) that several equally-valid post-hoc explanations could be put forward, we prefer to be cautious and simply report this effect without providing potentially incorrect interpretations.

We now turn to a discussion of the circuit potentially mediating the CMR, considering that we recorded LFPs from BA9 (Fig. 1), which is part of the dorso-lateral prefrontal cortex (dlPFC). This area (i.e. BA9), traditionally considered an high-order associative region, has been shown to also mediate motor functions, including the *control* of hand force, in both monkeys and humans (19–21). Specifically, an unilateral lesion of dlPFC (BA 9/10) in macaques impairs hand force control, leaving general motor behavior intact. Further, human studies have shown that this area is part of a network, including the basal ganglia, that subserves the control of grip force (19,20,43,44). In this context, the dlPFC has been suggested to be particularly important for real-time monitoring over the accuracy of force control, taking into account sensory feedback (43,44). These observations, taken together with our results, make dlPFC a suitable cortical region mediating the CMR. Notably, other RABs such as online motor correction or action stopping have been associated to activity of the dlPFC or to other prefrontal regions that are very proximal to dlPFC (1,16,45,46). This might potentially unify these distinct lines of research, and shed light upon a shared neural network mediating fast modulations of motor output in response to salient sensory stimuli (1). This issue deserves to be directly investigated in the future.

Through which pathway would BA9 influence the motor output and lead to the observed force modulations? Answering this question is not easy because there is no evidence of direct dlPFC projections to the spinal cord. Yet, we highlight three putative pathways that can indirectly influence the spinal cord. The first is the cortico-cortical route stemming from the supragranular layers (II-III) of BA9, which projects to the primary motor cortex through PMdr(F7) – PMdr(F2) – MI (47,48). The MI-corticospinal tract can convey force-related information to the spinal centers, through both slow and/or fast descending axons (49).

In addition, BA9 might reach the motor periphery by exploiting its massive projections to the paramedian, perpendicular, dorsomedial, and ventral pontine nuclei, as well as to the nucleus reticularis tegmenti pontis (50). This projection can be seen as the first feedforward segment of a prefrontal circuit reaching the cerebellum, which is implicated in different forms of adaptive behavior (51–53). At least in the cat, this circuit is made of both slow and fast-conducting axons (54) and is complemented by a feedback projection to prefrontal cortex from the cerebellar dentate nucleus (55). A third potential route is the dense bilateral area BA9 projection to the caudate nucleus, and the re-entrant signaling through the thalamus. This projection has recently been detailed in both monkeys and humans (56), and further elaborated in monkeys by Borra et al. (57). Which of these routes subserves the otherwise slow force response observed in our study, and their eventual interactions, remains to be determined.

Finally, it is important to highlight that recording from a single cortical area limits the interpretability of the results. This is particularly relevant given that sudden and unexpected stimuli activate large and widespread cortical territories, as suggested by the topographical distribution of the EEG vertex potential in both primates (Figure 3) and humans (5,58). We cannot therefore rule out the possibility that BA9 is not the specific site modulating the tonic motor output, and that other cortical areas would show similar pattern of LFP response and correlation with the CMR components. Studies entailing multiple intraparenchymal recordings will be necessary to test this likely alternative possibility.

### Conclusion

Our results provide compelling evidence indicating that salient sensory events modulate tonically-exerted isometric force following a CMR pattern. This behavioral response is both reactive and adaptive and is notably similar across human and non-human primates. The robust correlation between the positive wave of the vertex potential (measured with both EEG and LFP) and the force increase of the monkey CMR also matches what had been previously identified in EEG studies in humans. Notably, the LFP results provide strong evidence for a cortical origin of the CMR, and specifically highlight the role played by dlPFC, which can influence motor output by modulating distinct cortico-spinal and cortico-cerebellar pathways. As such, these results shed light upon a well-conserved cortical mechanism linking the detection of salient events to an immediate motor reaction, to prepare the animal for subsequent appropriate actions.

## Materials and Methods

### Animals and surgical procedures

Two male rhesus monkeys (*Macaca mulatta*) participated to the experiments: Monkey M (9 years old, 9.1 Kg) and Monkey T (9 years old, 9.4 Kg). One headpost was mounted on the skull in each animal. In between experiment 1 (EEG) and 2 (LFP), a circular chamber (diameter = 18 mm) was implanted for intracranial recording. The chamber was placed on the right hemisphere, centered at stereotaxic coordinates A +35; L +6 (Monkey 1) and A +35; L +7 (Monkey 2), in both cases corresponding to dorso-lateral prefrontal cortex (specifically BA 9) (see Supplementary Fig. 1). During the surgical procedures, the animals were pre-anaesthetized with ketamine (10 mg/kg, i.m.) and then anaesthetized with a mix of Oxygen/Isoflurane (1-3% to effect). Skull implants were performed under aseptic conditions. After surgery, the animals were allowed to recover for at least 7 days, while being treated with antibiotic and pain relievers, according to veterinary prescriptions. All efforts were made to minimize animals’ pain and distress. Animal care, housing and surgical procedures were in agreement with European (EU Directive 63-2010) and Italian (DL. 26/2014) laws on the use of non-human primates in scientific research.

### Experimental setup

The experimental setup is illustrated in Fig. 1a. Each monkey was placed in a soundproof chamber, seating on a primate chair in front of a 40-inch monitor (100 Hz, 800-600 resolution, 32-bit color depth; monitor-eye distance: 150 cm). Each animal was trained to control a colored circular cursor by applying a hand force on an isometric joystick [consisting of a 1.5-6.5-cm metal cylinder mounted on top of a force transducer: FTS-Gamma (Calibration SI-32-2.5) ATI Industrial Automation, Apex NC]. The cursor [0.6 degrees of visual angle (DVA)] was displayed on a black screen. The force exerted on the transducer was sampled (at 1 kHz) on both the x- and y-axes, corresponding to hand force exerted towards the left/right (x axis) and towards/away from the animal’s body (y axis) (Fig. 1a). Each animal faced the monitor from one out of two personalized primate chairs. Consequently, Monkey M had the monitor slightly on its right side (approximately 30 degrees from the midline, i.e. at 1 o’clock) while Monkey T slightly on its left-side (approximately at 11 o’clock).

The force exerted on the transducer was used to control the position of the cursor on the monitor, so that a force of 20N applied on the y axis (away from the animal’s body) was necessary to hold the cursor in the central target (Fig. 1). Sudden and unexpected auditory stimuli were produced using a beeper placed behind the monitor, ~160 cm away from the monkey’s head (Fig. 1a). Stimuli presentation and data sampling were controlled using the software package REX (59).

Both monkeys were required to use the left hand to perform the task, while the right arm was gently restrained. The joystick was controlled using the left hand because both monkeys appeared to prefer this configuration during the early stages of their training. We prevented the monkeys to reach their head with their arms by means of a 3D printed ‘safety box’ (designed using Autodesk Fusion 360), i.e. a nylon-12 surface that surrounded the animals’ neck and thus kept the EEG cap and electrodes away from the animals’ reach. Throughout the experiment, the monkey’s head was restrained using a titanium headpost.

### Behavioral task and paradigm

The task begun with the presentation of the target (an outlined grey circle, 2 DVA in diameter) placed in the center of the screen, together with the visual cursor (a white dot, 3 DVA diameter), placed below the target when no force was exerted on the isometric joystick (Fig. 1b). The monkey was required to bring the cursor inside the target, by exerting a force of 20N on the isometric joystick (on the y-axis, i.e. away from the body, Fig. 1a). The animal had to reach the target within 2 s from trial onset (i.e. presentation of the target), and keep the cursor within the target until the end of the trial (i.e. disappearance of the target). Trial duration ranged from 7 to 10 s (rectangular distribution). If the monkey did not reach the target within 2 s from its appearance or did not hold the cursor inside it for the whole trial duration, the trial was aborted. Otherwise, the trial was considered successful, and the animal received 1.75 ml of liquid reward (Fig. 1b).

### Experimental paradigm

While holding the cursor within the target, monkeys experienced two types of trials. On 1/3 of the trials, an auditory stimulus was presented (1 m distance, frequency 3.3 kHz, duration 50 ms). These trials are hereafter called “Beep trials”. The stimulus was always presented at least 3 s after the cursor had entered the target and not later than 3 seconds before the target disappearance. Within this time range, the timing of the stimulus was randomly assigned. On the remaining 2/3 of the trials, no auditory stimuli were presented, and monkeys were required to hold the cursor within the target for a comparable amount of time. These trials are hereafter called “No-Beep trials”. Beep and No-Beep trials were presented in a randomized order, within mini-blocks of 6 trials (2 Beep and 4 No-Beep trials), with the only caveat that no more than 2 Beep trials could be presented consecutively across successive mini-blocks.

### EEG equipment and montage (Experiment 1)

We recorded the electroencephalogram (EEG) using 29 active electrodes placed on the scalp (BioSemi Active-2 system). The data were sampled at 1024 Hz. The electrodes were mounted on two custom-made caps (http://www.easycap.de), tailored to fit each animal’s head, according to the layout displayed on fig. 1c.

The BioSemi system replaces the ground electrodes with two electrodes named CMS (Common Mode Sense, active electrode) and DRL (Driven Right Leg, passive electrode). According to the system’s guidelines, CMS should (ideally) be placed in the centre of the measuring electrodes, while DRL should be placed relatively away from them. While placing CMS, we also had to consider the position of the headpost, being approximately over Cz in monkey M, and over Cpz in monkey T. Therefore, CMS was placed on Cz (in monkey T) and on Cpz (in monkey M). DRL was always placed on frontal-left side of the animal’s head (see the layout displayed on Fig. 1c, CMS and DRL are highlighted using grey dots).

### Intracortical recordings (Experiment 2)

Neural raw signals were recorded from area BA9, using a 5-channel linear multiple-electrode array system for extracellular recording (Minimatrix 05. Thomas Recording, Germany). Inter-electrode distance was 0.3 mm. Each electrode (quartz-insulated platinum-tungsten fibers 80 mm diameter, 0.8–2.5 MOhm impedance) was guided through the intact dura into the cortical tissue (one specific recording site per session) through a remote controller. The raw neural signal was amplified, digitized at 24 kHz, and transmitted through optical fibers to a digital signal processing unit (RA16PA-RX5–2, Tucker-Davis Technologies) where it was stored.

### Data analysis (Experiment 1)

In Experiment 1 we collected 327 successful Beep trials for monkey M (12 recording sessions, 27.25 ± 16.33 trials per session) and 365 successful Beep trials for monkey T (8 recording sessions, 45.62 ± 8.44 trials per session). These data were analysed by applying the same pipeline (described hereafter) to the two datasets (one for each monkey) separately. This approach was preferred over the alternative ‘pooling’ over the two datasets (60) because the latencies of the force responses observed in the two animals were not always overlapping in time (see below).

#### Force analysis

Continuous force data were low-pass filtered (35 Hz, Butterworth, third order) and then segmented into epochs of 3 s. For Beep trials, the epochs started 0.4 s prior to stimulus onset and ended 2.6 s following it. For No-Beep trials, equally long epochs were extracted relatively to randomly-assigned time points comprised within the interval during which a stimulus could have been presented (i.e. at least 3 s after the cursor had entered into the target and not later than 3 s before the disappearance of the target). Force data comprised two channels F_x_ and F_y_ (associated with the force components exerted on the x and y axes of the transducer, respectively) and its magnitude F (which was computed using the following formula).

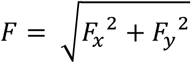

Trials contaminated by artifacts (i.e. deviating >4 SDs from the animal’s mean exerted force F across all trials) were excluded from further analyses (2,3). The corresponding EEG timeseries were also excluded. These trials constituted 3.01% (monkey T) and 4.28% (monkey M) of the total number of trials. Epochs were baseline corrected using the −0.05 to 0 s prestimulus interval (2,3). Beep and No-Beep trials were compared using two-sample t-tests (one for each timepoint).

#### EEG analysis

Continuous EEG data were band-pass filtered (1-35 Hz, Butterworth, third order) and then segmented into Beep and No-Beep epochs of 5 s (−1.4 s to 3.6 s). Because the datasets contained several movement artifacts, data pre-processing was assisted by a validated algorithm for automatic artifact-correction: Artifact Subspace Reconstruction (ASR, threshold value = 5) (36,37). ASR is an adaptive algorithm based on principal component analysis. It estimates clean portions of data to determine thresholds that are later used to reject large-variance components. The use of ASR was preferred over conventional ‘data cleaning’ procedures because of its automaticity, implying lower computational time and lesser (potentially arbitrary) decision-making. We note that we also compared the current results to those obtained following a traditional ‘data cleaning’ procedure, which yielded similar results at the cost of several trials being rejected.

Following ASR, the EEG epochs were cropped to match the force epochs (i.e. −0.4 to 2.6 s). Noisy or faulty electrodes were interpolated by replacing their voltage with the average voltage of the neighbouring electrodes. Data were re-referenced using a common average reference (61). Artifacts due to eye blinks or eye movements were subtracted using a validated method based on an independent component analysis (62). In all datasets, independent components related to eye movements had a frontal scalp distribution. We also estimated the voltage at electrodes Cz and Cpz (used for CMS and for the headholder) by computing the average voltage of the neighbouring electrodes. Finally, the EEG epochs were baseline corrected using the −0.2 to 0 s prestimulus interval. Beep and No-Beep trials were compared using paired-sampled t-tests (one for each timepoint).

The trial-by-trial correlation between EEG and force magnitude (F) epochs was computed consistently with our previous work (2,3). Specifically, we first smoothed the signals using a moving average (sliding window = 20 ms). The signals were then resampled to 250 Hz to reduce computation time. Finally, the trial-by-trial correlation coefficient (Spearman’s r) was computed between EEG amplitude and force magnitude, for all possible pairs of time points between the interval −50 to 400 ms of the EEG time course (i.e., the interval encompassing all EEG modulations) and the interval −50 to 2000 ms of the force time course (i.e., the interval encompassing all force modulations). This resulted in 29 correlation matrixes (one for each EEG electrode). Significant correlations were thresholded by extracting clusters encompassing at least two consecutive significant timepoints (p<0.05) associated to at least two neighbouring electrodes.

### Data analysis (Experiment 2)

In Experiment 2 we collected 393 successful Beep trials for monkey M (28 recording sessions, 14.04 ± 4.05 trials per session) and 339 successful Beep trials for monkey T (25 recording sessions, 13.56 ± 1.90 trials per session).

Behavioural data from Experiment 2 were analysed by applying the same pipeline described for Experiment 1. Trials contaminated by artifacts (i.e. deviating >4 SDs from the animal’s mean exerted force F across all trials) were excluded from further analyses. The corresponding LFP timeseries were also excluded. These trials constituted 3.20% (monkey M) and 4.78% (monkey T) of the total number of trials.

Continuous extracellular LFP data were band-pass filtered (1-35 Hz, Butterworth, third order), polarity-inverted (for comparability with the EEG signal), and then segmented into Beep and No-Beep epochs of 5 s (−1.4 s to 3.6 s). LFP data were recorded from 5 electrodes, each with a single recording site. Within each recording session, a variable number of electrodes failed to penetrate the dura and did not reach the target cortical depth. These electrodes were considered ‘non-active’, and their corresponding LFP timeseries were excluded from the analyses [69 out of 140 (49.29%) for monkey M, and 15 out 125 (12.00%) for monkey T]. The remaining ‘active’ electrodes were classified as ‘superficial’ or ‘deep’ by applying a median split on the cortical depth from which recordings were taken.

The trial-by-trial correlation between LFP and force epochs was computed as in Experiment 1. Correlation matrixes were calculated by pooling all ‘active’ electrodes together or by pooling ‘superficial’ or ‘deep’ electrodes separately. Significant correlations were thresholded for significant time intervals (p < 0.05).

---

**Supplementary Figure 1.**
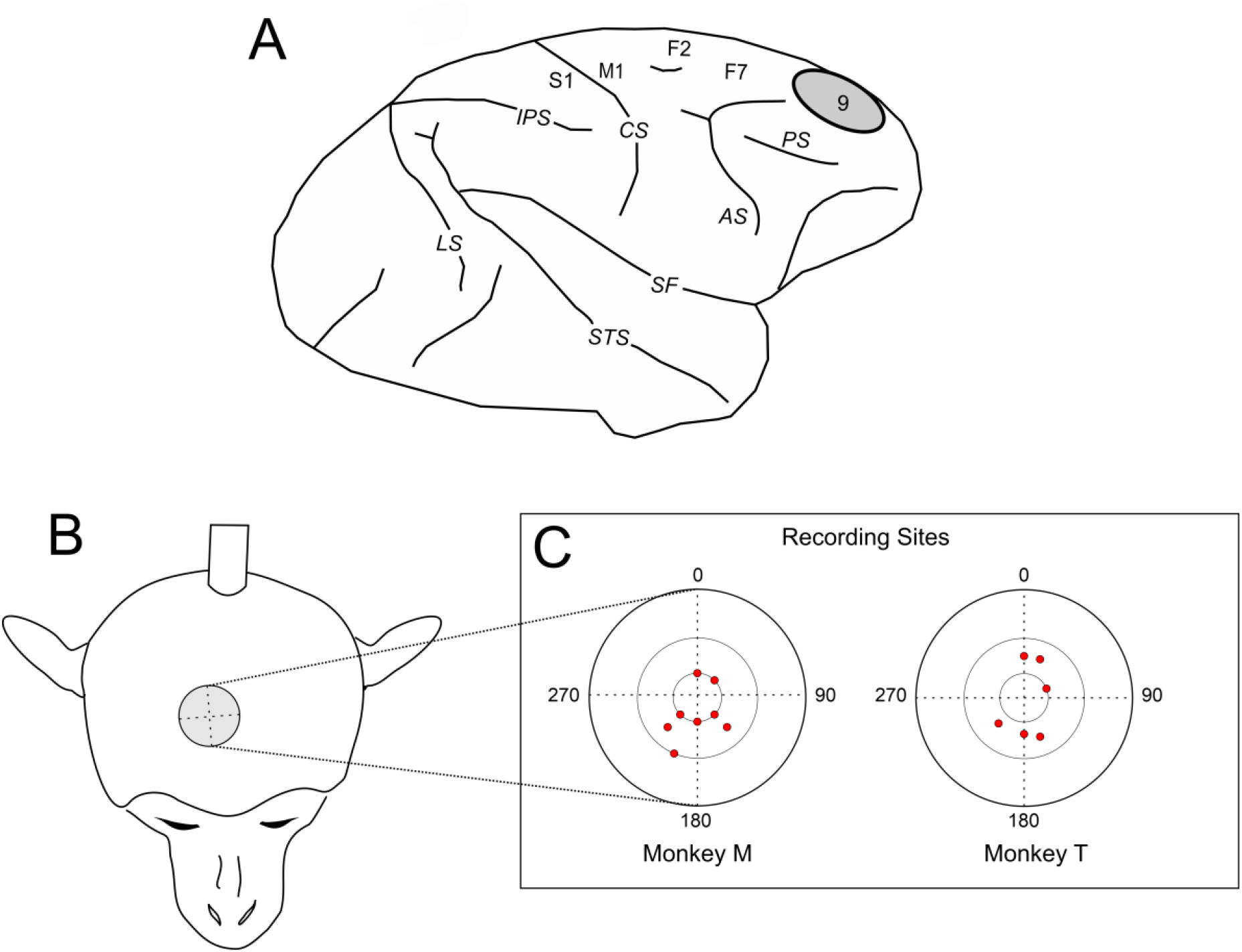
Recordings of Local Field Potentials. **(A)** Schematic representation of the cortical site (BA9) from which Local Field Potentials were recorded (AS Arcuate Sulcus, PS Principal Sulcus, CS Central Sulcus, IPS Intraparietal Sulcus, LS Lunate Sulcus, STS Superior Temporal Sulcus, SF Sylvian Fissure). **(B)** The chamber (diameter = 18 mm) was placed on the right hemisphere, centered at stereotaxic coordinates A +35; L +6 (Monkey 1) and A +35; L +7 (Monkey 2). **(C)** Detailed location of the recording sites, reported separately for the two monkeys. Each site (red dots) included up to five microelectrode penetrations.

**Supplementary Figure 2.**
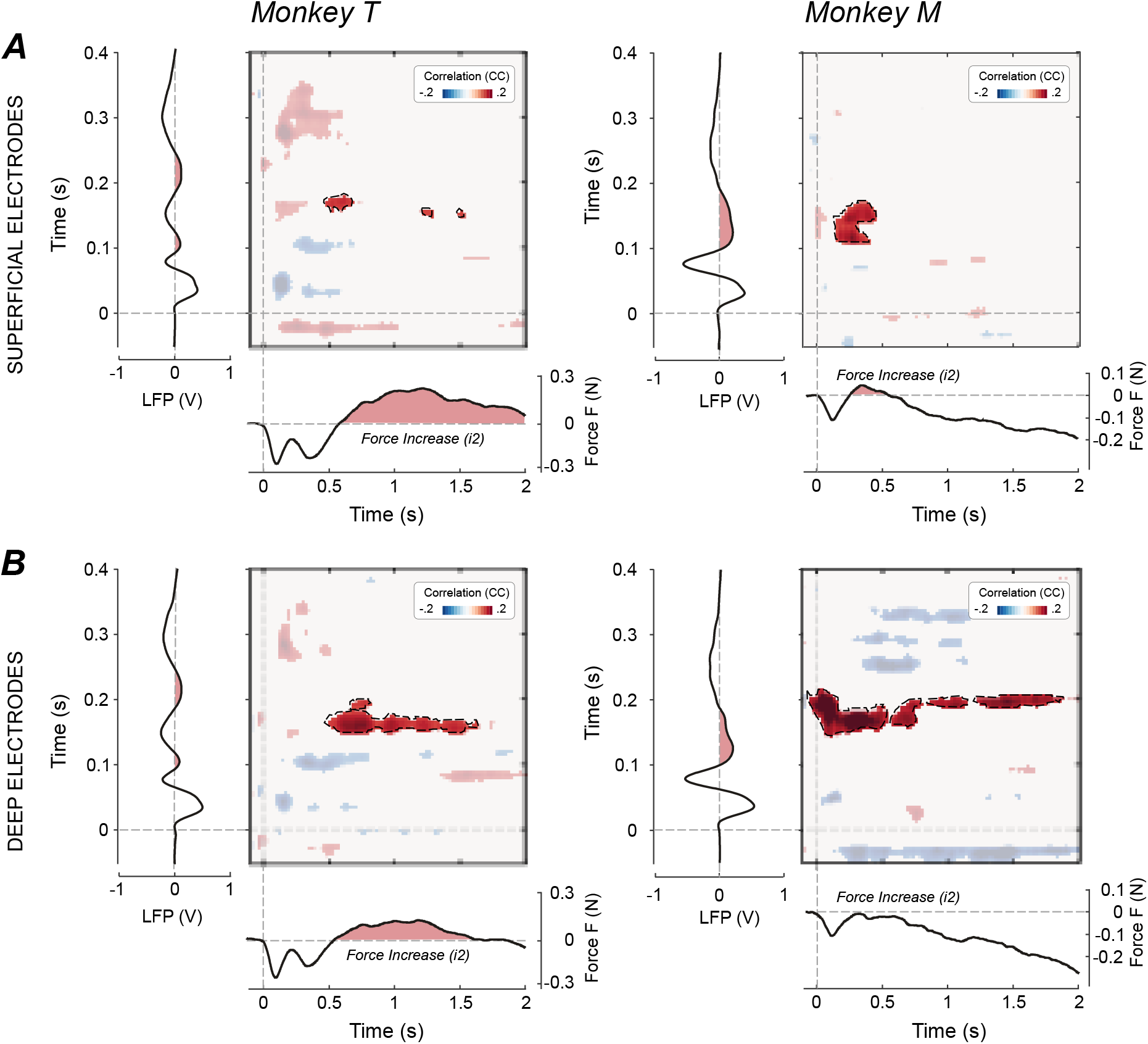
Trial-by-trial correlations between stimulus-induced force and LFP modulations at different cortical depths. The bidimensional plots represent the significant trial-by-trial correlation coefficients (cluster-corrected Spearman’s r) between LFP and force for all possible pairs of time points, for either superficial (A) or deep (B) recording sites. The LFP timeseries (plotted vertically) and the force timeseries (plotted horizontally) are shown to assist interpretability of the correlations. The correlation between the LFP positive wave (equivalent to the EEG P130) and the force increase (i2) is highlighted. Note the clearer LFP-force correlations in deep recording sites.

## Acknowledgments

GN acknowledges the support of the European Research Council (Starting Grant MUSICOM). GDI acknowledges the support of the European Research Council (Consolidator Grant PAINSTRAT) and of The Wellcome Trust (COLL JLARAXR). ABM acknowledges the support of The Italian Ministry of Education (Grant. N. 201794KEER_002).

